# The dynamic neural code of the retina for natural scenes

**DOI:** 10.1101/340943

**Authors:** Niru Maheswaranathan, Lane T. McIntosh, Hidenori Tanaka, Satchel Grant, David B. Kastner, Josh B. Melander, Aran Nayebi, Luke Brezovec, Julia Wang, Surya Ganguli, Stephen A. Baccus

## Abstract

Understanding how the visual system encodes natural scenes is a fundamental goal of sensory neuroscience. We show here that a three-layer network model predicts the retinal response to natural scenes with an accuracy nearing the fundamental limits of predictability. The model’s internal structure is interpretable, in that model units are highly correlated with interneurons recorded separately and not used to fit the model. We further show the ethological relevance to natural visual processing of a diverse set of phenomena of complex motion encoding, adaptation and predictive coding. Our analysis uncovers a fast timescale of visual processing that is inaccessible directly from experimental data, showing unexpectedly that ganglion cells signal in distinct modes by rapidly (< 0.1 s) switching their selectivity for direction of motion, orientation, location and the sign of intensity. A new approach that decomposes ganglion cell responses into the contribution of interneurons reveals how the latent effects of parallel retinal circuits generate the response to any possible stimulus. These results reveal extremely flexible and rapid dynamics of the retinal code for natural visual stimuli, explaining the need for a large set of interneuron pathways to generate the dynamic neural code for natural scenes.

---

Nearly all of our understanding of retinal computations and circuit mechanisms comes from artificial stimuli such as flashing spots, drifting gratings and white noise ^1-3^, which have unknown relevance to natural visual processing. Although numerous retinal computations have been identified by such methods including selectivity for specific types of motion, adaptation to various statistics and prediction of visual features^3^, the number of interneurons (>40) is even greater, suggesting an undiscovered complexity in retinal processing. To characterize the neural code for natural scenes, given that the vertebrate retina has three layers of cell bodies, we tested whether three layer convolutional neural network (CNN) models (Figure 1) could predict the responses of populations of salamander retinal ganglion cells responding to a 50-minute sequence of either natural images or spatiotemporal white noise. Natural scene images changed every second, and were jittered with the statistics of fixational eye movements^4,5^, creating a spatiotemporal stimulus. The model had eight different model cell types in each of the first and second layers that tiled the visual field, with each cell type having a distinct receptive field, and a final layer that represented the responses of individual ganglion cells. We found that CNN models could predict the responses of ganglion cells to either natural scenes or white noise nearly up to a fundamental limit of precision set by intrinsic neural variability, and were substantially more accurate than linear-nonlinear (LN) models^6^ or generalized linear models (GLMs) ^1^ (Figure 1B, C). Based on results varying the number of cell types in the first two layers, eight cell types were chosen as the minimum number that achieved the maximal model performance (Fig. 1D).

**Figure 1.**
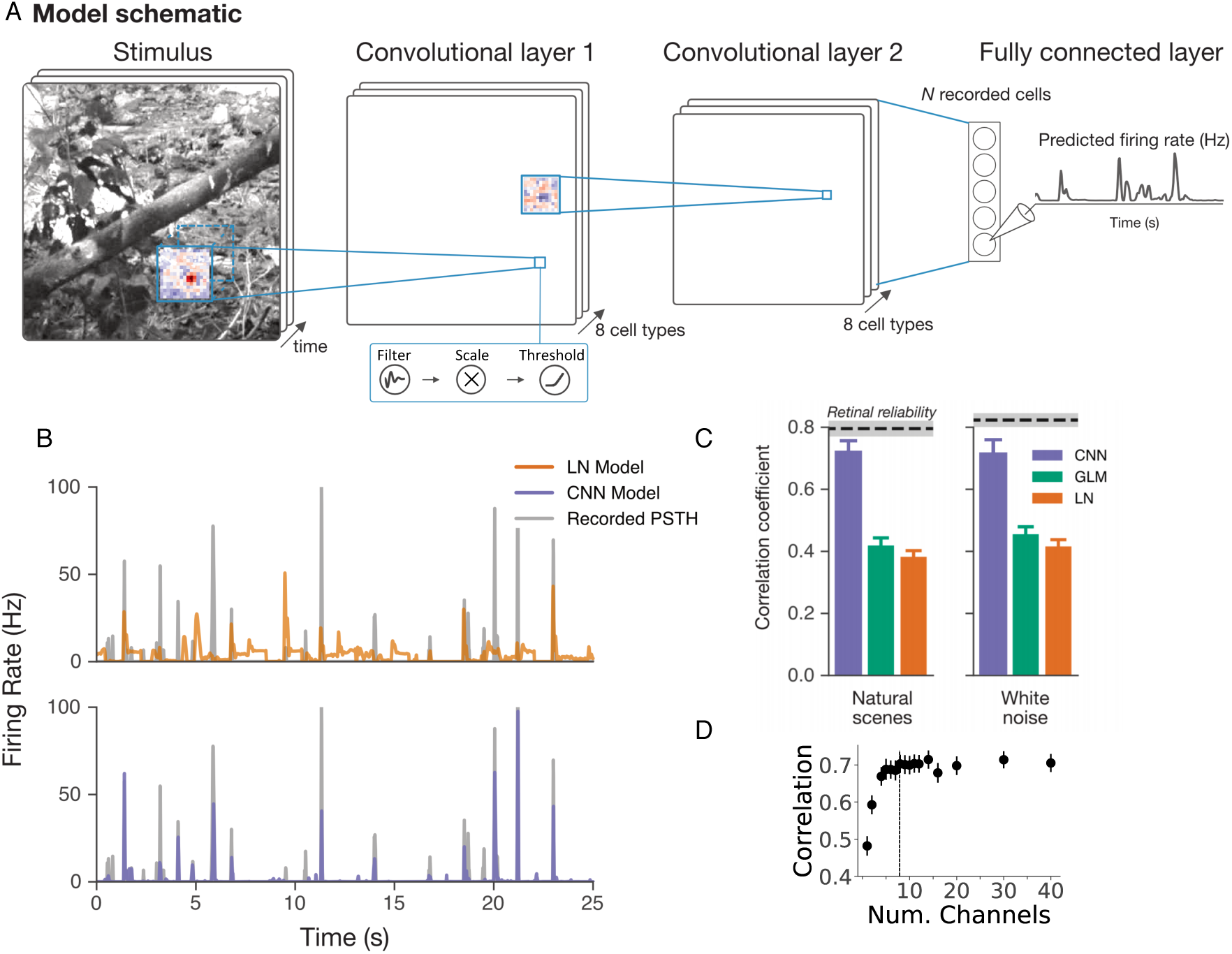
Convolutional neural networks provide accurate models of the retinal response to natural scenes. (A) Convolutional neural network model trained to predict the firing rate of simultaneously recorded retinal ganglion cells from the spatiotemporal movie of natural scenes. The first layer is a spatiotemporal convolution, the second is a spatial convolution, and the third is a final dense layer, with rectifying nonlinearities in between each layer. Each location within the model also has a single parameter that scales the amplitude of the response. (B) PSTHs comparing recorded data and Linear-Nonlinear (LN) or CNN models for the test data set. (C) Comparison of LN, Generalized Linear Model (GLM) and CNN model predictions for a 25 second segment of a natural scene movie. Correlation coefficients are for the test data set, as compared to the retinal reliability of ganglion cell PSTHs correlated between different sets of trials (dotted line is mean, grey bar is 1 s.e.m.) (D) Correlation coefficient between model and test data set for different numbers of cell types in the first and second layer. Dashed line indicates 8 cell types, the value chosen for further analysis in this study.

## CNNs internal units are highly correlated with interneuron responses

To examine whether the internal computations of CNN models were similar to those expected in the retina, we computed receptive fields for first and second layer model cell types in CNNs trained on responses to natural scenes. We found that the receptive fields of CNN model cells had the well-known structure of retinal interneurons^7,8^, with a spatially localized center-surround structure (Fig. 2A-B), a mix of On and Off responses, and both monophasic and biphasic temporal filters.

**Figure 2.**
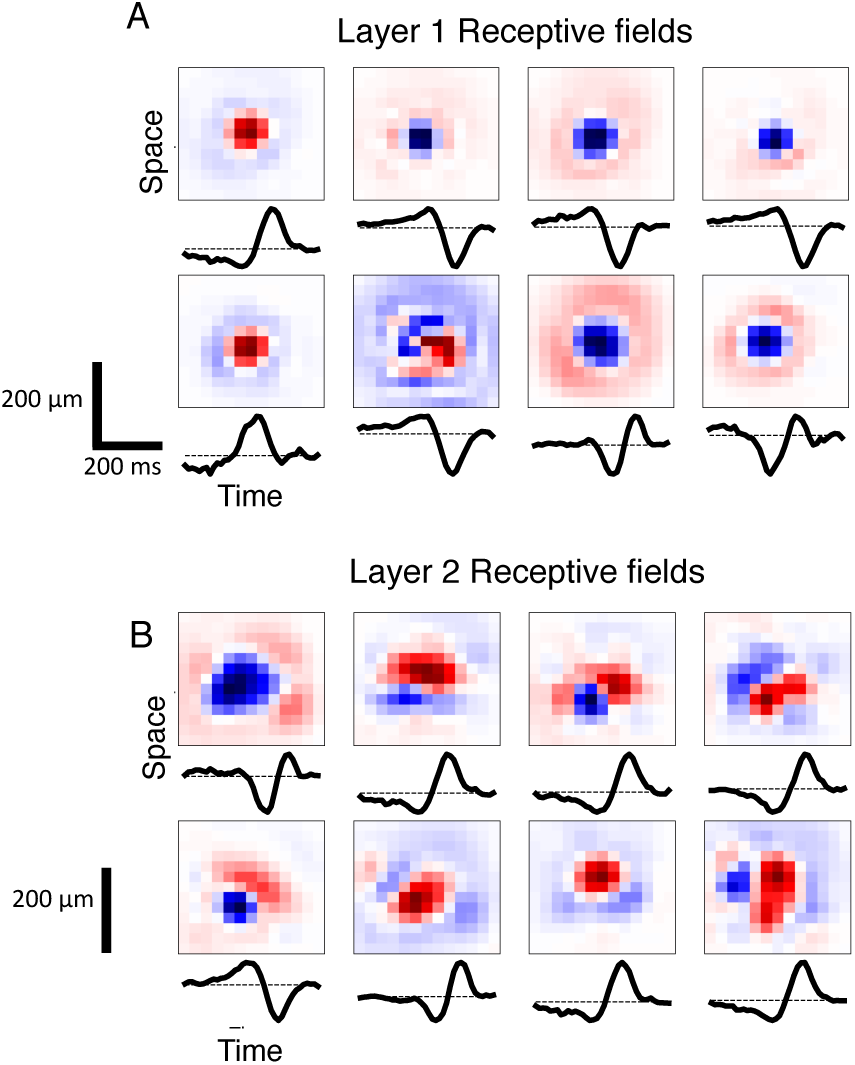
Structure of receptive fields of model cell types. (A) Receptive fields of model units in Layer 1 computed by presenting a white noise stimulus to the model, and shown as the spatial and temporal average of the space-time separable approximation to the receptive field. (B) Same for Layer 2.

In inferotemporal cortex, units of CNNs have been shown to be correlated with a linear combination of the activity of individual neurons^9^ making it difficult to draw conclusions about individual neurons by an examination of CNN units. We compared the activity of CNN units to interneuron recordings performed on separate retinae that the model was never fit to (Fig. 3A). The stimulus presented to the retina and separately to the model was a spatiotemporal white noise checkerboard, a stimulus that has no spatiotemporal correlations except for the 50 µm size and 10 ms duration of square stimulus regions. We compared each interneuron recording with 8 units of the first layer and 8 units of the second layer at each location to find the most correlated unit in the model at the location of the cell. We found that each recorded interneuron was highly correlated with a particular unit type, and only at a single location (Figure 3B-E). Spatiotemporal receptive fields were highly similar between recorded interneurons, and their most correlated model cell type (Fig. 3B). The magnitude of this correlation approached the variability of the interneurons themselves, as assessed by using an LN model fit to the interneuron to predict another segment of the interneuron’s own response (Figure 3 C,D). This correlation was specific for individual unit types, as could be observed by ranking the model cell types from most to least correlated, and finding that the second and lower most correlated model cell types were substantially less correlated with the interneuron than the most correlated cell type (Fig. 3 D,E). This high correlation did not arise by chance as a “null” CNN model fit by shuffling spikes relative to the stimulus did not produce internal units correlated with interneuron responses (Fig. 3D, E). Therefore, fitting a CNN model to the natural scene responses of retinal ganglion cells alone models an entire population of interneurons, many of which have high correlation with measured interneuron responses created with a different stimulus and a different retina.

**Figure 3.**
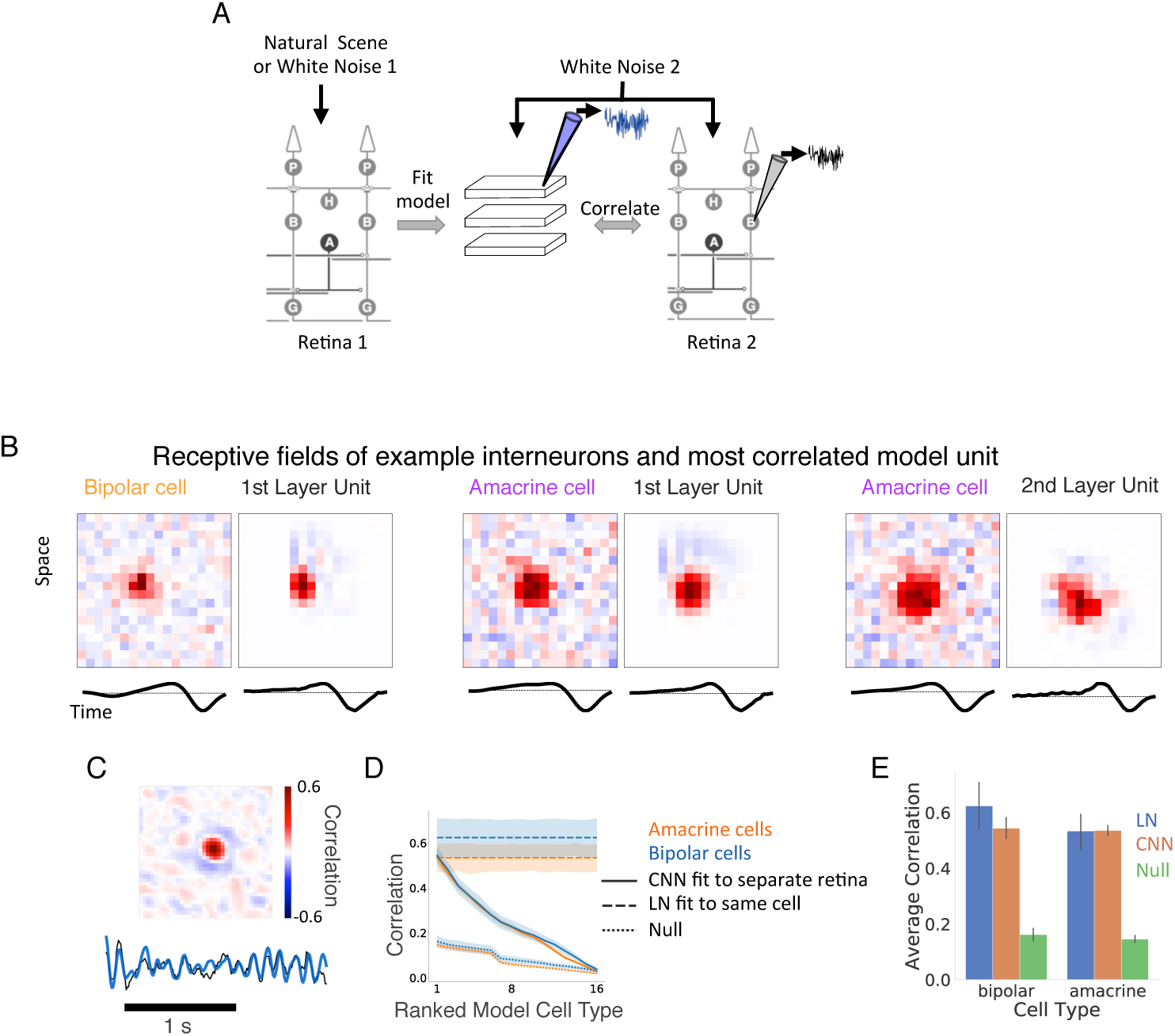
Model internal units are correlated with interneuron responses. (A) Schematic of experiment. Models were fit to natural scenes or white noise stimuli. Bipolar or amacrine cells from a different retina were recorded intracellularly responding to a different white noise sequence. (B) Spatiotemporal receptive fields of example interneurons recorded from a separate retina, and the model unit that was most correlated with that interneuron. The model was never fit to the interneuron’s response. (C) Top. Correlation map of a model cell type with the response of an interneuron recorded from a different retina to a white noise stimulus. Each pixel is the correlation between the interneuron and a different spatial location within a single model cell type. Bottom. Responses compared to the most correlated model unit and the interneuron. (D) The average correlation between different interneuron types (7 bipolar, 26 amacrine) and model cell types ranked from most correlated model unit (left) to least (right). Dashed lines indicate the correlation between an interneuron’s response and an LN model fit to a separate segment of the recording from the same interneuron. Thus, the correlation between model units and interneurons approaches the variability of the interneurons themselves. Dotted lines indicates correlation between interneuron responses and a null model, fit after taking the spikes of a ganglion cell in 5 second blocks and shifting them randomly relative to the stimulus (E). Average correlation between interneuron recordings and the most correlated CNN unit from a different retina, an LN model fit to the same interneuron, and the null model.

## A wide range of retinal phenomena are engaged by natural stimuli

Numerous nonlinear computations have been identified by presenting artificial stimuli to the retina, including flashing spots, moving bars and white noise. However we neither understand to what degree natural vision engages these diverse retinal computations elicited by artificial stimuli, nor understand the relationship between these computations under natural scenes and underlying retinal circuitry. We tested models fit either only to natural scenes or white noise by exposing them to a battery of structured stimuli previously used in the literature to identify and describe retinal phenomena. We focused on effects shorter than 400 ms, which was the longest timescale our model could reproduce as limited by the first layer spatiotemporal filter. Remarkably, the CNN model exhibited fast contrast ^10-12^ adaptation (Fig. 4A), latency encoding ^3^ (Fig. 4B), synchronized responses to motion reversal ^13^(Fig. 4C), motion anticipation ^14^ (Fig. 4D), the omitted stimulus response ^15^ (Fig. 4E), frequency doubling in response to reversing gratings ^16^ (Fig. 4F) and polarity reversal ^17^ (Fig. 4G). All of these response properties arose in a single CNN model simply as a by-product of optimizing the models to capture ganglion cell responses to natural scenes. CNN models trained on white noise did not exhibit all of these phenomena, in particular failing to capture fast contrast adaptation, latency encoding and the omitted stimulus response, indicating that natural scene statistics trigger nonlinear computations that white noise does not. Even though these natural scenes consisted only of a sequence of images jittered with the statistics of fixational eye movements (the stimulus contained no explicit object motion or periodic patterns), the CNNs still exhibited motion anticipation and reversal, and the omitted stimulus response.

**Figure 4.**
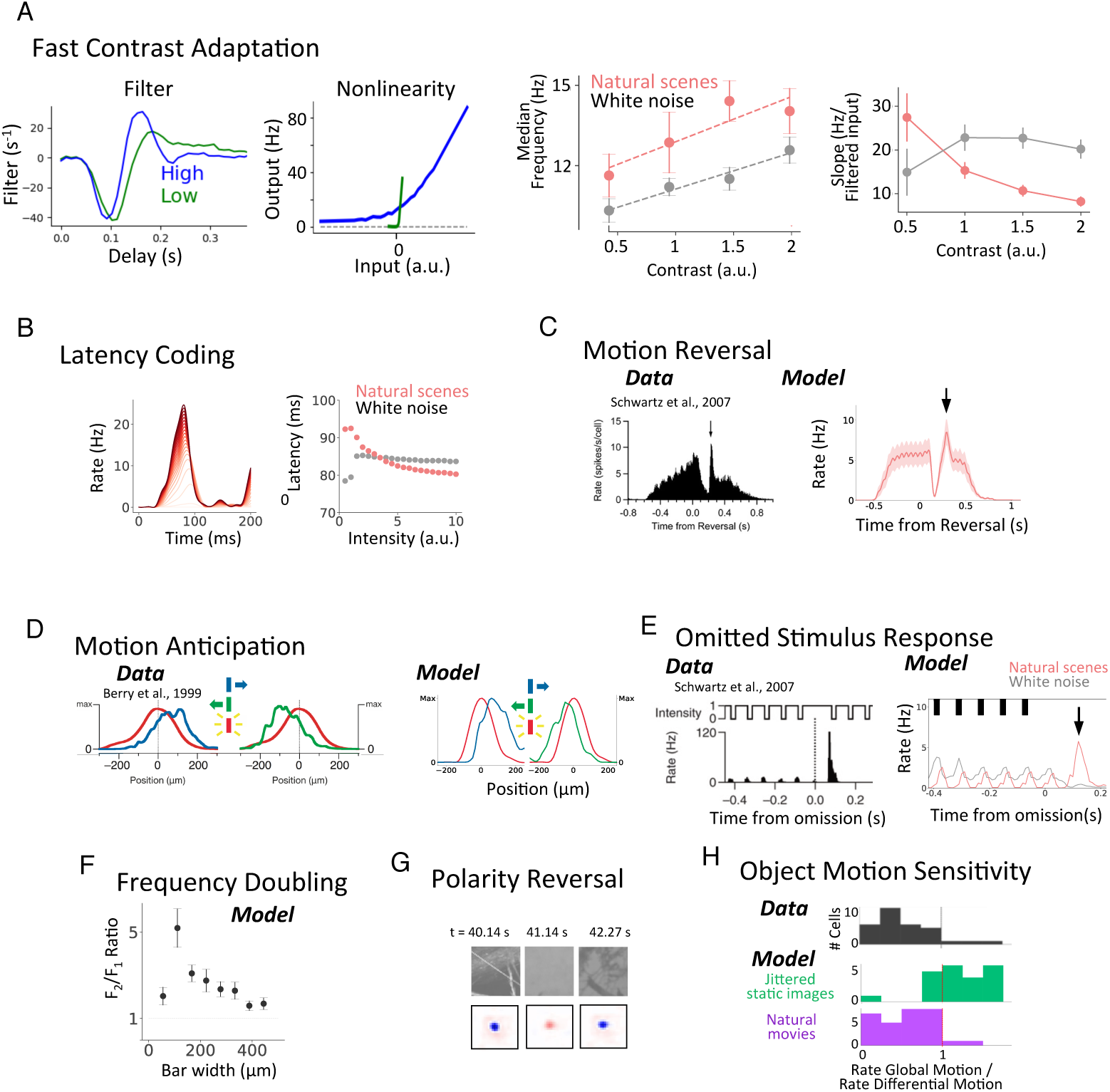
CNN models reveal that many nonlinear retinal computations are engaged in natural scenes. After fitting a model to natural scenes, a number of artificially structured stimuli were presented to the model. (A) Contrast adaptation. Left. LN model of a ganglion cell responding to a uniform field stimulus with low or high contrast, showing adaptive changes in temporal filtering and gain. Middle, Median temporal frequency taken from the Fourier transform of the temporal filter, averaged over a population of ganglion cells as a function of contrast. Results shown for models fit to natural scenes and white noise. Right, Averaged gain measured as the slope of the nonlinearity as a function of contrast, showing that CNN models decrease their decrease their gain with contrast when fit to natural scenes, but not when fit to white noise. (B) Latency encoding. Left: Flash response with intensities ranging from weak to strong. Right: Latency of the peak response vs. stimulus intensity for models trained on natural scenes or white noise. (C) Motion reversal. Stimulus consists of a moving bar that abruptly reverses direction at different positions. Left. Published results of a population of ganglion cells showing a synchronous response (arrow) to the reversal. Also shown is the population response of CNN model cells. (D) Motion anticipation. Population ganglion cell responses to a flashed bar (red) vs motion to the right (blue) or left (green), from published results^18^ (left) or the CNN model (right). (E) Omitted stimulus response (OSR). Left. Published results^19^ showing the response to a missing stimulus following a train of flashes. Right. CNN model response to a sequence of three flashes. The OSR (arrow) appears for models trained on natural scenes but not white noise. (F) Frequency doubling in response to reversing gratings of different width, computed as the ratio of the response at twice the stimulus frequency (F2) and the response at the stimulus frequency (F1). (G) Polarity reversal. Example reversal of polarity during a natural image sequence. Each panel shows the current image (top) and corresponding instantaneous receptive field (bottom) for an example cell at a fixed delay (∼100 ms) relative to the stimulus at different times during the sequence, showing fast kernel reversal from an OFF-feature (blue) to ON-(red) and back. Rapid receptive field changes are further analyzed in Fig. 5. (H). Object Motion Sensitivity. CNN models were fit to either jittered static images or natural movies consisting of swimming fish in the presence of image jitter and saccade-like transitions. Stimuli were then shown to the model consisting of a jittering central grating surrounded by a jittering background grating. Gratings moved either synchronously (Global motion) representing eye movements, or asynchronously (Differential Motion) representing object motion. Shown is the ratio of firing rates in Global Motion to Differential Motion. A ratio much less than one indicates Object Motion Sensitivity. Results for (A-F) are from a population of 26 ganglion cells. Figures reproduced with permission from authors.

The only retinal phenomenon tested that was not captured by the model was the object motion sensitive (OMS) response^5^, a computation thought to discriminate object motion from retinal motion due to eye movements. We hypothesized that the absence of an OMS response in the model was due to the lack of differential motion in the training stimulus, and trained additional models on the retinal response to movies of swimming fish that include differential motion. We found that these models did indeed exhibit an OMS response (Fig. 4H). Thus the model reveals whether retinal computations triggered by one stimulus occur in another, in particular during natural scenes.

## Interneuron contributions to a dynamic visual code

Receptive fields in sensory neuroscience are typically thought of as representing a static sensory feature, although it is known that this feature can change slowly due to adaptation to the statistics of the stimulus ^18-20^. A particularly advantageous property of CNN models is that rapid dynamics of visual sensitivity can be examined by computing the instantaneous receptive field (IRF), which can be easily calculated as the gradient of the model output with respect to the current stimulus (Fig. 5A). This can be done at each moment of time, allowing us to examine for the first time the full dynamics of the receptive field and assign those dynamics to the action of interneurons.

**Figure 5.**
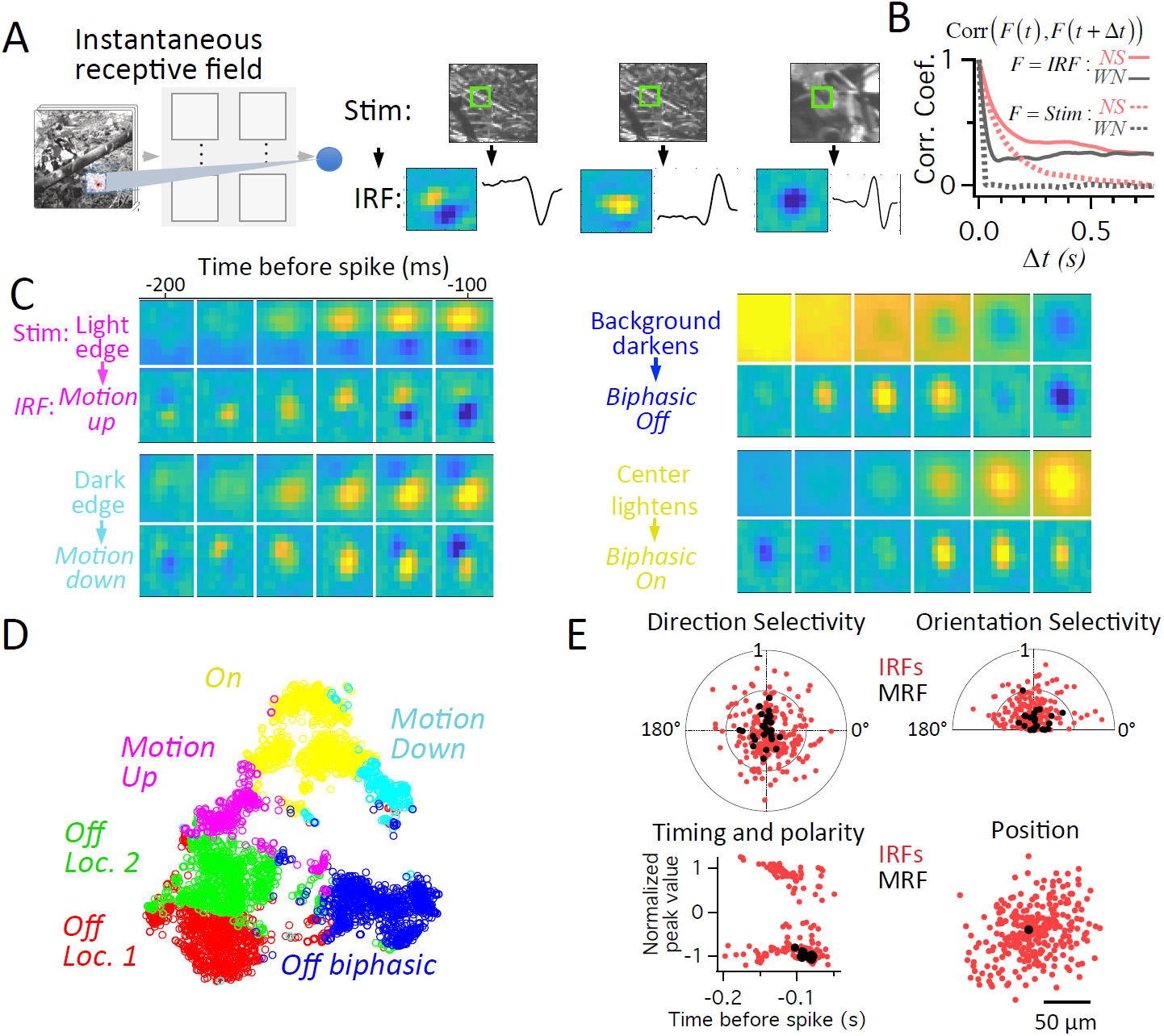
Dynamic mode switching of retinal receptive fields. (A) Diagram of the instantaneous receptive field (IRF) as the sensitivity of the ganglion cell to the stimulus at each moment. (B) Average correlation coefficient between IRFs at different times separated by a time interval Δ*t* for white noise and natural scenes. Also shown for comparison are average correlations between stimulus frames. (C) Four IRF clusters for a single cell. Top Row: Average spatiotemporal stimulus that drove an IRF cluster, shown as a sequence of stimulus frames from 200 ms to 100 ms preceding a spike. Bottom Row: The mean spatiotemporal IRF in each cluster. Left, Two different IRF clusters showing motion sensitivity (Top: motion up, Bottom: motion down), which were driven by an edge. Right. IRF clusters driven by intensity changes showing a biphasic OFF receptive field when the background intensity changed, and a biphasic On receptive field when the stimulus center brightened. (D) t-sne analysis of IRFs, colored by cluster identity from k-means clustering of IRFs performed separately. (E) Top left. For 26 neurons with 12 IRFs clusters each, radial axis shows Direction Selectivity Index for IRFs and mean receptive field (MRF), plotted against the preferred angle of motion. Top right. Same for Orientation Selectivity Index. Bottom left. Normalized value of the first peak (either positive or negative) of the temporal filter, plotted against the time of the peak for IRFs and MRF. Bottom right. The position of the center of mass of IRFs relative to the center of mass of the MRF (black point at zero). 50 µm corresponds to ∼ one visual degree.

IRFs changed with extremely rapid dynamics on the scale of tens of ms, as judged by comparing the correlation coefficient between IRFs at different time delays. The dynamics of the IRF were limited by stimulus correlations, in that for an uncorrelated stimulus (white noise), the IRF changed from its previous value with a time constant of ∼ 30 ms (Fig 5B). To examine these rapid changes in feature selectivity, we clustered the IRFs computed at each time point, revealing that during natural stimuli the retina signaled in different modes that changed rapidly depending on the stimulus frame (Fig. 5, C - E). By computing the average stimulus for each IRF cluster, we unexpectedly found new phenomena triggered by different ethologically relevant stimuli. The presence of an edge changed the IRF to be maximally sensitive to motion of that edge, and the direction of motion and orientation preference changed with the intensity gradient of the edge (Fig 5 C, E). IRFs also showed much stronger direction and orientation selectivity than could be observed in the mean receptive field (MRF) (Fig 5E). The location of the IRF showed also substantial variation compared to the MRF (Fig. 5E). Furthermore, changes in local stimulus intensity reversed the polarity of the IRF (Fig. 5E), indicating a local effect distinct from previously reported polarity reversal triggered by peripheral stimuli^17^.

Because the IRF is a mathematical sum of the features conveyed by different interneuron pathways in the model, we investigated the source of these dynamic receptive fields by performing an exact decomposition of a ganglion cell’s response into the Interneuron Contributions (INCs) of each of the 8 model cell types in the first layer at each time point (Fig. 6), using the method of Integrated Gradients ^21,22^ (see methods). Intuitively, the INC is the product of the sensitivity of the model interneuron to the stimulus and the sensitivity of the model output to the model interneuron, and is an application of the multivariable chain rule. Thus to determine the effect of an interneuron on the circuit’s output, this analysis takes account of both a cell’s input (its receptive field) and its output (projective field) ^23^. To assess the model interneuron’s contribution over a range of stimuli rather than a single point in stimulus space, the INC is integrated over a straight path of increasing contrast from the zero stimulus, a grey screen, to the particular stimulus frame. This new type of analysis is different than simply examining the representation of a stimulus in a neural population, and reveals how an interneuron population uses that stimulus representation to change the model circuit’s output.

**Figure 6.**
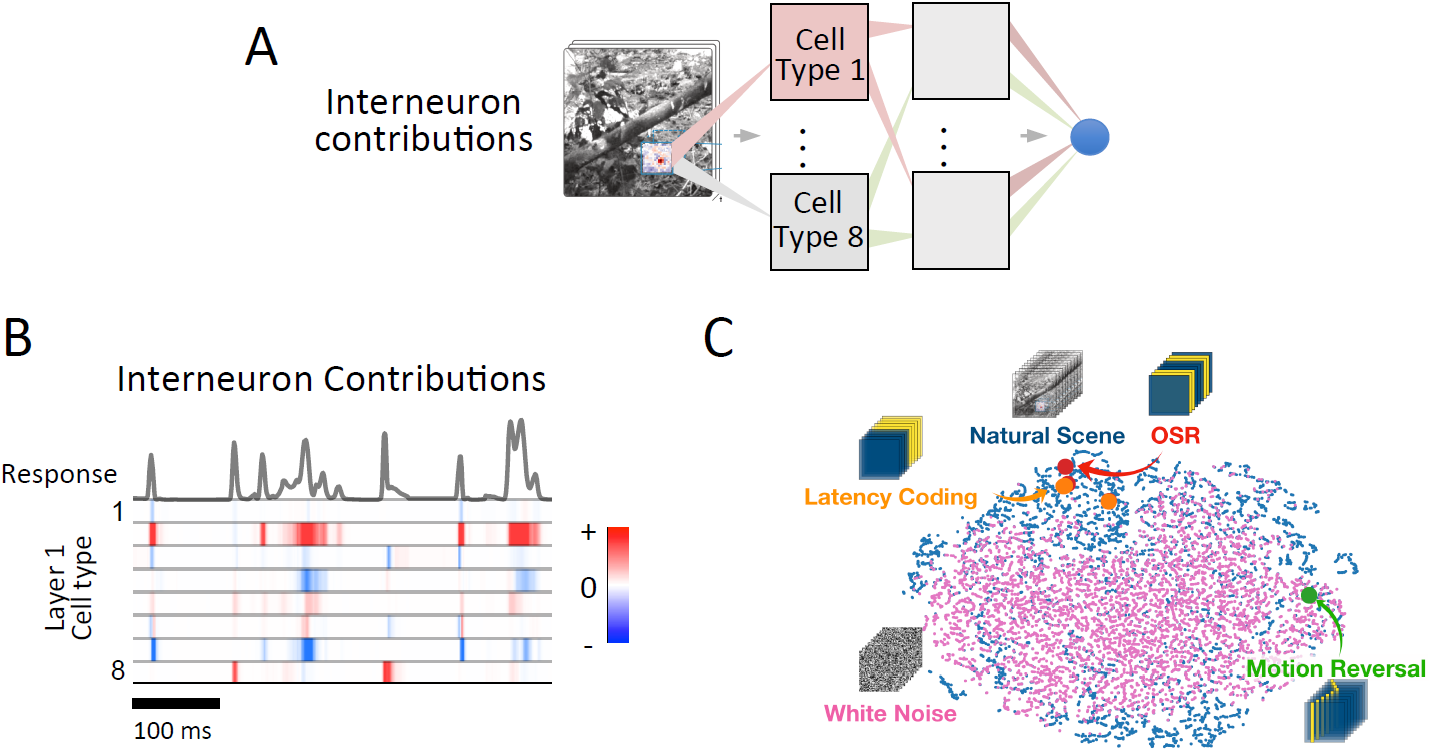
Interneuron contributions to natural and artificial scenes. (A) Diagram of concept of Interneuron Contributions (INCs), which represent how much each model unit (cell) contributes to the model’s output for each particular stimulus (see methods). We focused on the contribution of Layer 1 model units, and averaged over all units of a given type (B) INCs for the 8 cell types for the model’s first layer for a natural stimulus sequence. Each colored row shows the contribution of a cell type in layer 1 of the model. (C) t-SNE plot including natural scenes, white noise, and several artificially structured stimuli that can be summarized by a single 400 ms stimulus sequence. Each point in the plot represents the vector of interneurons in layer 1 contributing to the response at a single time point.

We identified the patterns of INCs that generate the code for natural stimuli, white noise and artificially structured stimuli. We found surprisingly that the interneuron patterns generating responses to some artificial stimuli live within the space of those elicited by natural stimuli but not within the space of white noise (Fig. 6 B, C), showing that these artificially structured stimuli are indeed ethologically relevant to understanding the retinal code under natural scenes. This further explains why models fit to natural scenes but not white noise recapitulated the previously described phenomena triggered by these structured stimuli – white noise is insufficient to explore the stimulus space that triggers these phenomena, but natural scenes are. Thus, natural scenes drive the set of interneuron contributions into a set of states that encompasses previously explored artificial stimuli, showing the relevance of stimuli of unknown functional relevance such as the omitted stimulus response^15^.

These results capture the dynamic retinal code of natural scenes, and connect that code to much of the retinal phenomenology previously described. This approach reveals the extensive rapid changes of the neural code on a previously inaccessible timescale, and enables a direct determination of the contribution of cell types to any arbitrary stimulus. Because model cell types have high correlation with retinal interneurons, this approach will serve as the foundation to define how interneuron patterns generate the dynamic neural code for natural scenes.

## Methods

### Visual Stimuli

A video monitor projected the visual stimuli at 30 Hz controlled by Matlab (Mathworks), using Psychophysics Toolbox ^24^. Stimuli had a constant mean intensity of 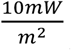. Images were presented in a 50 × 50 grid with a square size of 25 µm at a frame rate of 100 Hz. Static natural jittered scenes consisted of images drawn from a natural image database ^25^ and drifted in two dimensions with the approximate statistics of fixational eye movements^5^. The image also changed to a different location every one second, representing a saccade-like transition. Natural movies consisted of fish swimming in an aquarium, and contained both drift and saccade-like transitions that matched static jittered natural scenes. For analysis of model responses to artificial stimuli (Fig. 3), unless otherwise stated stimuli were chosen to match published values for each phenomenon.

### Electrophysiology

Retinal ganglion cells of larval tiger salamanders of either sex were recorded using an array of 60 electrodes (Multichannel Systems) as previously described ^26^. Intracellular recordings were performed using sharp as previously described.

### Model training

We trained convolutional neural network models to predict retinal ganglion cell responses to either a white noise or natural scenes stimulus, simultaneously for all cells in the recorded population of a given retina ^27^. Model parameters were optimized to minimize a loss function corresponding to the negative log-likelihood under Poisson spike generation,

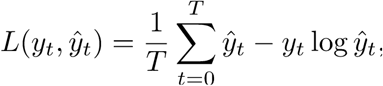

where 𝒴*t* and 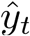 are the actual and predicted firing rates of the retinal ganglion cells at time *t*, respectively with a batch size of *T*, chosen to be 50 s. To help with model fitting, we smoothed retinal ganglion responses during training with a 10 ms standard deviation Gaussian, the size of a single time bin in our model.

The architecture of the convolutional neural network model consisted of three layers, with 8 cell types (or channels, in the language of neural networks) per layer. Each layer consisted of a linear spatiotemporal filter, followed by a rectification using a rectified linear unit (ReLU). For each unit, an additional parameter scaled the activation of the model unit prior to the rectified nonlinearity. This scaling parameter could vary independently with location.

Optimization was performed using Adam ^28^, a variant of stochastic gradient descent. Models were trained using TensorFlow^29^ or PyTorch^30^ on NVIDIA Titan X GPUs. Training an individual model to convergence required ∼8 hours on a single GPU. The networks were regularized with an L2 weight penalty at each layer and an L1 activity penalty at the final layer, which helped maintain a baseline firing rate near 0 Hz.

During optimization, the spatial components of linear filters were implemented as a series of stacked linear convolutions, each consisting of a series of 3 × 3 filters. Thus seven sequential 3 × 3 filters were applied to generate a 15 × 15 filter. After optimization, these sequential filters were collapsed into a single linear filter. Therefore, this procedure did not change the final architecture of the model, but improved the model’s performance, presumably by reducing the number of parameters.

We split our dataset into training, validation, and test sets, and chose the number of layers, number of filters per layer, the type of layer (convolutional or fully connected), size of filters, regularization hyperparameters, and learning rate based on performance on the validation set. We found that increasing the number of layers beyond three did not improve performance, and we settled on eight filter types in both the first and second layers, with filters that were much larger (Layer 1,15 × 15 and Layer 2, 11 × 11) compared to traditional deep learning networks used for image classification (usually 5 × 5 or smaller). Values quoted are mean s.e.m. unless otherwise stated.

#### Linear-Nonlinear Models

Linear-nonlinear models were fit by the standard method of reverse correlation to a white noise stimulus ^6^. We found that these were highly susceptible to overfitting the training dataset, and imposed an additional regularization procedure of zeroing out the stimulus outside of a 500 µm window centered on the cell’s receptive field.

#### Generalized Linear Models

Generalized linear models (GLMs) were fit by minimizing the same objective as used for the CNN, the Poisson log-likelihood of data under the model. We performed the same cutout regularization procedure of only keeping the stimulus within a 500 µm region around the receptive field (this was critical for performance). The GLMs differed from the linear-nonlinear models in that they have an additional spike history feedback term used to predict the cell’s response (Pillow et. al. 2008). Instead of the standard exponential nonlinearity, we found that using soft rectified functions log(1+exp(x)) gave better performance.

### Interneuron contributions

To identify the contribution of each model neuron to the processing of specific visual stimuli, we used the recently developed method of Integrated Gradients ^21,22^ to decompose a ganglion cell’s firing rate into the Interneuron Contributions (INCs) of each of the 8 model cell types by performing path integral. Mathematically, the trained deep learning model represents a nonlinear function *r*(*t*) = 𝓕[*s*(*t*)], where *r*(*t*) is the output firing rate and *s*(*t*) ∈ *R*^50×50×40^ is the movie input. Using the line integral *s*(*t*; *α*) = *αs*(*t*) where the path takes a straight line *α* : 0 → 1, and assuming 0 = 𝓕[*s*(*t*; 0)] we obtain an equality

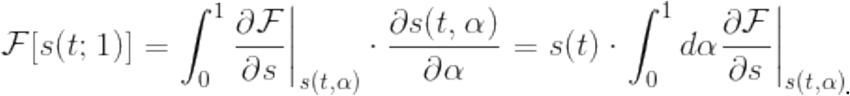

Our goal is to quantify the contributions of the first layer model units 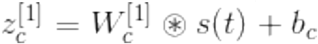, where “[1]” refers to an index of the layer, “*c”* refers to channel, 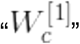 is the linear convolutional filter, and *b*_c_ is the bias parameter. Therefore we further apply the chain rule to define the INC of *c*-th channel (*A*_c_) as

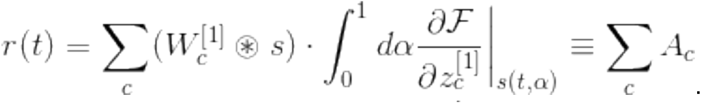

Finally, the spatially averaged INCs *A*_c:1∼8_ forms a vector with eight elements, which is taken as the contribution of that model cell type to the model output at that instant of time.

## Acknowledgments

The authors wish to acknowledge William Newsome, Jennifer Raymond, Tom Clandinin and Leonidas Guibas for helpful discussions. This work was supported by grants from the NEI, Pew Charitable Trusts, McKnight Endowment Fund for Neuroscience and the Ziegler Foundation, (SAB); Burroughs Wellcome, McKnight, James S. McDonnnell, Simons Foundations, and the Office of Naval Research (SG), an NSF fellowship (NM) an NRSA (LTM), by the Stanford Medical Scientist Training Program (DBK) and an NSF IGERT graduate fellowship (DBK),

